# An Unexpectedly Complex Architecture for Skin Pigmentation in Africans

**DOI:** 10.1101/200139

**Authors:** Alicia R Martin, Meng Lin, Julie M Granka, Justin W Myrick, Xiaomin Liu, Alexandra Sockell, Elizabeth G. Atkinson, Cedric J Werely, Marlo Möller, Manjinder S Sandhu, David M. Kingsley, Eileen G Hoal, Xiao Liu, Mark J. Daly, Marcus W Feldman, Christopher R Gignoux, Carlos D Bustamante, Brenna M Henn

**Affiliations:** Department of Genetics, Stanford University, Stanford, CA, 94305; Analytic and Translational Genetics Unit, Department of Medicine, Massachusetts General Hospital and Harvard Medical School, Boston, MA 02114; Program in Medical and Population Genetics, Broad Institute, Cambridge, MA 02141; Stanley Center for Psychiatric Research, Broad Institute, Cambridge, MA 02141; Department of Ecology and Evolution, SUNY Stony Brook, NY 11794; Department of Biological Sciences, Stanford University, Stanford, CA, 94305; BGI-Shenzhen, Shenzhen, China; Department of Developmental Biology, Stanford University, Stanford, California, USA; SA MRC Centre for Tuberculosis Research, DST/NRF Centre of Excellence for Biomedical Tuberculosis Research, Division of Molecular Biology and Human Genetics, Faculty of Medicine and Health Sciences, Stellenbosch University, Tygerberg, South Africa; Wellcome Trust Sanger Institute, Genome Campus, Hinxton, England.

**Keywords:** pigmentation, human evolution, Africa, heritability, population genetics

## Abstract

Fewer than 15 genes have been directly associated with skin pigmentation variation in humans, leading to its characterization as a relatively simple trait. However, by assembling a global survey of quantitative skin pigmentation phenotypes, we demonstrate that pigmentation is more complex than previously assumed with genetic architecture varying by latitude. We investigate polygenicity in the Khoe and the San, populations indigenous to southern Africa, who have considerably lighter skin than equatorial Africans. We demonstrate that skin pigmentation is highly heritable, but that known pigmentation loci explain only a small fraction of the variance. Rather, baseline skin pigmentation is a complex, polygenic trait in the KhoeSan. Despite this, we identify canonical and non-canonical skin pigmentation loci, including near *SLC24A5*, *TYRP1*, *SMARCA2*/*VLDLR*, and *SNX13* using a genome-wide association approach complemented by targeted resequencing. By considering diverse, under-studied African populations, we show how the architecture of skin pigmentation can vary across humans subject to different local evolutionary pressures.

**Highlights:**

- Skin pigmentation in Africans is far more polygenic than light skin pigmentation in Eurasians.
- KhoeSan^§^ populations, which diverged early in human prehistory from other populations, have lightened skin pigmentation compared to equatorial Africans.
- Skin color is highly heritable in the KhoeSan, but pigmentation variability is not well explained by previously discovered pigmentation genes.
- We perform the first GWAS for pigmentation in African KhoeSan populations and identify canonical pigmentation loci near *TYRP1* and in *SLC24A5*, as well as novel associations surrounding *SMARCA2* and other genes.

## Introduction

Skin pigmentation is one of the most strikingly variable and strongly selected phenotypes among human populations (Sabeti et al., 2007; Sturm and Duffy, 2012), with darker skin observed closer to the equator and lighter pigmentation observed at high latitudes. Researchers have hypothesized that variable exposure to ultra violet radiation (UVR) creates opposing selective forces for vitamin D production and folate protection, resulting in global pigmentation differentiation (Chaplin and Jablonski, 2009; Jablonski and Chaplin, 2010). Skin pigmentation differences at similar latitudes and UV exposures indicate that additional evolutionary forces, such as assortative mating, drift, and epistasis, are also likely to have affected global skin pigmentation (Wilde et al., 2014; Pośpiech et al., 2014). While ~171 genes have been implicated in pigmentation variability across model organisms (e.g. the Color Genes database: http://www.espcr.org/micemut/), only ~15 genes have been associated with skin color differences in humans (**Table 2**). The relative paucity of skin pigmentation loci identified from GWAS efforts has led to the characterization of pigmentation variation as a relatively simple trait, with only a handful of SNPs being highly predictive of skin, eye, and hair color across populations (Hart et al., 2013; Spichenok et al., 2011; Walsh et al., 2013).

The genetic basis of skin pigmentation has been primarily studied in Europeans, Asians, and admixed individuals of western African descent using candidate gene and genome-wide approaches (Candille et al., 2012; Beleza et al., 2013a; Beleza et al., 2013b; Sulem et al., 2007; Sulem et al., 2008; Sturm and Duffy, 2012). Remarkably, only one study of quantitative genetic effects on pigmentation in continental Africans has been published to date, despite the fact that Africans have the greatest range of pigmentation variation globally (Relethford, 2000; Jablonski and Chaplin, 2014; Crawford et al, 2017). Several adaptive sweeps have occurred at large-effect skin pigmentation loci in populations from high latitudes, which researchers have interpreted as resulting from strong environmental selection pressure that reduces variability in the population. For example, *SLC24A5* is among the best-studied skin pigmentation genes; the derived Ala111Thr allele (rs1426654) confers the largest known lightening effect and has swept to fixation in western Eurasian populations (Beleza et al., 2013a; Lamason et al., 2005). rs35395 near *SLC45A2*, rs10831496 near *GRM5* and *TYR*, and rs4424881 near *APBA2* and *OCA2* also have different allele frequencies in Europeans and Africans, with high derived frequencies that confer large skin lightening effects in Europeans (Beleza et al., 2013a; Norton et al., 2007). Other variants of smaller effect contribute to the relatively narrow variation among Europeans in skin pigmentation, including associations in/near *MC1R*, *TYR*, *IRF4*, and *ASIP* (Sulem et al., 2007; Sulem et al., 2008). Several of the largest skin lightening effects in Europeans and East Asians arose through convergent evolution; for example the His615Arg amino acid substitution in *OCA2* (rs1800414) has a large functional skin lightening effect specifically in East Asians (Yang et al., 2016). Some of the same genes have been selected across Eurasia at high latitudes with evidence of similar (e.g. *KITLG*) (Miller et al., 2007) or different (*OCA2*) selective sweeps, whereas other genes have been selected/associated with skin pigmentation in populations living close together at high latitudes, such as *ATRN* and *DCT* in East Asians (McEvoy et al., 2006; Lao et al., 2007; Edwards et al., 2010) or *SLC45A2*, *SLC24A5*, and *TYRP1* in Europeans (Soejima et al., 2006; Soejima and Koda, 2007; Izagirre et al., 2006; Voight et al., 2006; Lao et al., 2007).

Strong positive selection acting on skin pigmentation has resulted in large effect sizes that explain a large fraction of heritable variation. For example, a previous study showed that only 4 loci explain 35% of the variation in skin pigmentation in recently admixed Cape Verdeans, who have European and West African ancestors (Beleza et al., 2013b). In contrast, complex traits such as height and schizophrenia, typically require ~10,000 independent SNPs derived from GWAS of >100,000 individuals to build predictors that respectively explain ~29% and ~20% of the variance in independent cohorts (Wood et al., 2014; Schizophrenia Working Group of the Psychiatric Genomics Consortium, 2014). As a consequence of strong positive selection, previous studies of naturally selected traits such as pigmentation, high altitude adaptation, and response to pathogens have repeatedly shown that these traits have typically evolved substantially larger effect sizes than, for example, complex common disease; these large effect loci have typically been discovered with small sample sizes (i.e. ~100s of individuals) relative to GWAS of complex, highly polygenic traits (Kenny et al., 2012; Yi et al., 2010; Zhou et al., 2013; Alkorta-Aranburu et al., 2012; Genovese et al., 2010; Kayser et al., 2008; Moltke and Albrechtsen, 2013). While adaptive pigmentation loci are among the most diverged in the genome across populations, it is worth nothing that effect size estimates for nearly all significant GWAS associations that are polymorphic across well-studied populations are on average directionally consistent at individual loci across populations (Liu et al., 2015; Carlson et al., 2013), but that, in aggregate, prediction accuracy varies across populations (Martin et al., 2017).

Populations at lower latitudes—closer to the equator—typically have darker skin color than those in Europe or East Asia. Recently admixed populations (i.e. groups with highly divergent ancestors) have increased pigmentation variation, and the largest genetic effects are due to derived alleles that reduce melanin (Marcheco-Teruel et al., 2014; Beleza et al., 2013b; Norton et al., 2006; Norton et al., 2007). For example, all four significantly associated pigmentation loci identified in the study of Cape Verdeans were derived in Europeans, which together with admixture proportions explain the majority of heritable variation (Beleza et al., 2013b). Together, these studies highlight a complex interaction between latitude, the strength of selection (i.e. likelihood of selective sweep), and the distribution of effect sizes (i.e. polygenicity). A clear understanding of the genetic determinants of dark skin variability is lacking.

Striking skin pigmentation variability among African populations has been underappreciated in genetic studies (Relethford, 2000; Jablonski and Chaplin, 2014). Additionally, genetic diversity declines with distance from Africa (Ramachandran et al., 2005), which, coupled with greater phenotypic variation, suggests that more genetic variation may contribute to skin pigmentation diversity in Africa. Light skin pigmentation is observed in the far southern latitudes of Africa among KhoeSan hunter-gatherers and pastoralists of the Kalahari Desert and nearby regions. The KhoeSan are unique in their early divergence from other populations, likely dating back at least ~100,000 years ago (Gronau et al., 2011; Schlebusch et al., 2012; Veeramah et al., 2012); they exhibit extraordinary levels of genetic diversity and low levels of linkage disequilibrium (LD) (Henn et al., 2011; Li et al., 2008; Ramachandran et al., 2005). Previous work points to southern Africa as the point of origin for modern humans (Henn et al., 2011; Luca et al., 2011; Tishkoff et al., 2009), but it is unknown whether moderate to light skin pigmentation in the different KhoeSan groups is an example of convergent evolution with northern Europeans and Asians, or reflects the ancestral human phenotype. Previous studies using samples from the Human Genome Diversity Panel (HGDP) have noted different pigmentation allele frequencies between the Ju|’hoansi KhoeSan and other Africans, but these have been based on n<7 individuals from the former population without associated phenotype data (Berg and Coop, 2014; Norton et al., 2007).

Here we report an evolutionary and genetic study of skin pigmentation with a total of 465 genotyped KhoeSan individuals (278 ‡Khomani San and 187 Nama), with targeted resequencing at associated pigmentation loci in 439 KhoeSan individuals (268 ‡Khomani, 171 Nama) and matched quantitative spectrophotometric phenotype data (**Table S4**). The ‡Khomani San are traditionally a N|u-speaking hunter-gatherer population living in the southern Kalahari Desert, while the Nama are traditionally a Khoekhoe-speaking semi-nomadic pastoralist group of KhoeSan ancestry. We investigate: i) the degree of polygenicity and heritability of skin pigmentation, ii) the extent to which variation in pigmentation is explained by previously associated or canonical pigmentation genes, and iii) novel pigmentation alleles contributing to variation in the ‡Khomani San and Nama populations.

## Results

Baseline skin color was quantitatively phenotyped for 479 individuals (277 ‡Khomani, 202 Nama, **Table S4**) via specialized narrow-band reflectometry that specifically measures hemoglobin and melanin of both the left and right upper inner arms (Methods, (Diffey et al., 1984)), with M index = log_10_(1 / % red reflectance). Sequencing and/or genotyping was performed for a subset of phenotyped samples (**Table S4**, Methods). Skin pigmentation is considerably lighter in the KhoeSan than the majority of other African populations, with baseline upper arm M index = 57.57 ± 10.12 (mean ± sd, N = 278) in the ‡Khomani San. Baseline upper arm pigmentation in the Nama is slightly lower, with M index = 52.12 ± 8.93 (N=223). The ‡Khomani are on average significantly darker than the Nama (p=3.6e-10, Figure 1C), but the variance is not significantly different (p>0.05). For comparison, we aggregated quantitative skin pigmentation across 32 globally diverse populations (4,712 individuals) assayed with a DermaSpectrometer (DSMI or DSMII—the latter was used in this study) (Basu Mallick et al., 2013; Beleza et al., 2013b; Candille et al., 2012; Durazo-Arvizu et al., 2014; Edwards et al., 2010; Norton et al., 2006; Coussens et al., 2015) (Figure 1A-B, **Table S1**). Only four African populations are available for comparison; among these only the Ghanians represent an equatorial African population without recent admixture. Skin color is substantially darker in equatorial Ghanaians, where M index reaches a mean of 96.04 ± 10.94; M index for Cape Verdeans, who have ~40% European admixture on average, have slightly lighter (55.39 ± 13.00, p=5.6e-3) and considerably more variable pigmentation (p=1.9e-6) than the KhoeSan. Two other populations living in South Africa, the Xhosa and admixed Coloured populations, have respectively darker (M index=67.1±7.5) and similar (M index=53.1±8.5) pigmentation compared to the KhoeSan populations (Coussens et al., 2015).

**Figure 1.**
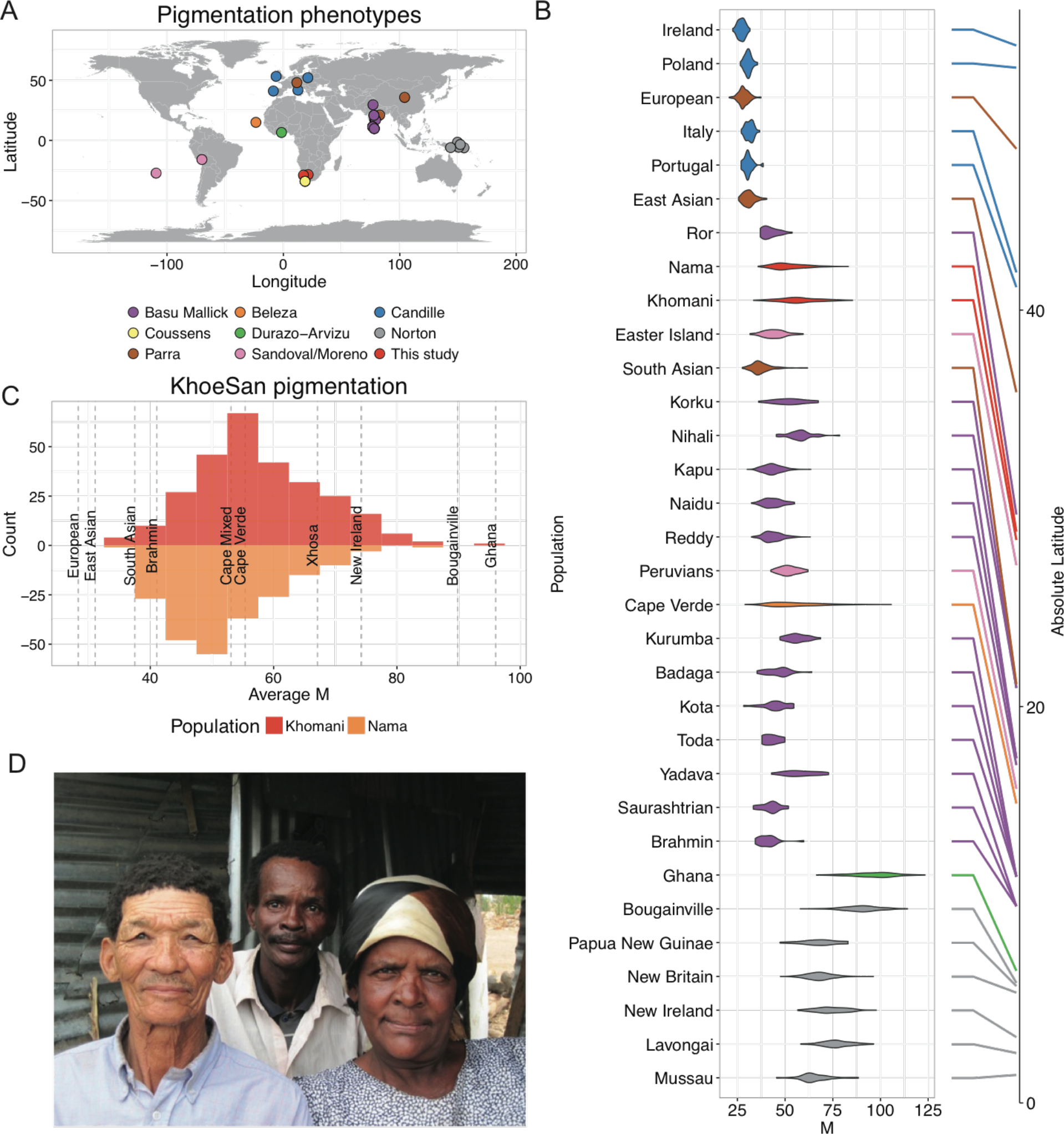
Distributions of baseline pigmentation in globally diverse populations. **A**) Sample locations of skin pigmentation datasets where phenotypes were measured with a DermaSpectrometer I or DermaSpectrometer II. **B**) Violin plots of pigmentation distributions for 32 populations from 8 studies ordered by latitude; absolute latitudes provided on the right. Corresponding datasets are colored as in A). **Table S1** provides summary statistics for each population. M indices are reflectance measures that approximate melanin content. **C**) A comparison of skin pigmentation distributions in ‡Khomani (top) and Nama populations (bottom). Dashed grey lines and labels indicate mean M index for the indicated other global populations. **D**) South African individuals in a household that exemplify the substantial skin Absolute Latitude pigmentation variability in the ‡Khomani and Nama populations. Picture taken with consent for publication.

### Evidence of Increased Polygenicity in Skin Pigmentation Among Equatorial Populations

We tested whether the correlation between absolute latitude and pigmentation was significant with our large, quantitatively phenotyped sample of global populations. As previously observed (Jablonski and Chaplin, 2010; Byard, 1981; Zaidi et al., 2017), we find that skin pigmentation is strongly associated with absolute latitude (R^2^=0.53, β=−1.18 on M index scale, p<2e-16); populations further from the equator have lighter skin pigmentation. We next tested whether variance in melanin within populations also varies across populations. Skin pigmentation has primarily been studied in lightly pigmented European and East Asian populations, where skin color varies minimally among individuals (Figure 1A-B). Less-studied equatorial and admixed populations, including Melanesians, Ghanaians, Cape Verdeans, South African admixed Coloured, and South Asians vary considerably more in skin pigmentation (Figure 1B). We find that absolute latitude is also significantly negatively associated with the standard deviation in melanin (R^2^=0.41, p=5.0e-5). Further, melanin distributions are heteroskedastic (i.e. the variance is not constant—rather, it changes over the range of observed M index), with the coefficient of variation, a standardized metric of phenotypic dispersion, decreasing with increasing distance from the equator (cv=σ/μ, R^2^=0.14, p=0.03, **Table S1**).

A sign test comparing variances in lighter versus darker population pairs within the same study indicates that populations with lighter skin have significantly reduced phenotypic variance than expected by chance (p=2.01e-8). These results suggest that there is reduced genetic heterogeneity and/or reduced variance in the population distribution of causal effect sizes contributing to lighter versus darker pigmentation. There is more than an order of magnitude difference in variance between the lightest and darkest populations (i.e. Irish vs Ghanaian F=0.03, p=6.7e-23). Europeans and East Asians have significantly less variation than South Asians (F=0.25, p=1.06e-14 and F=0.30, p=1.27e-10, respectively, Figure 1B). Cape Verdeans with the highest quartile of European admixture have lighter, less variable skin color than individuals with the lowest quartile of European ancestry (p=4.28e-9, although notably ancestry proportions are bimodal across individuals). Among Melanesians, islands at similar latitudes with more lightly pigmented individuals on average show less variance than those with more darkly pigmented individuals (e.g. one-sided F test comparing variance among more lightly pigmented New Britain individuals versus individuals from Bougainville, p=2.89e-9, Figure 1B). Among the ‡Khomani and Nama, comparing individuals with primarily European admixture (>20%, N=124) to individuals with primarily Bantu admixture (>20%, N=91), we find significantly greater melanin variation among KhoeSan individuals with more Bantu admixture (p=1.33e-4).

### Ancestry and Skin Pigmentation Variation in the KhoeSan

The ‡Khomani San and the Nama have both experienced admixture with neighboring darker-skinned Bantu-speaking groups beginning ~450 years ago, as well as with lighter-skinned European settlers who first arrived in the Northern Cape during the late 18^th^ century (Uren et al., 2016). We assessed these ancestry proportions using unsupervised allele frequency clustering with ADMIXTURE as well as principal components analysis (PCA, Methods). At *k*=3, we observe distinct clustering between Europeans, Bantu-speaking and West African populations, and KhoeSan populations; both the Nama and the ‡Khomani have ~75-80% KhoeSan-specific ancestry. For *k*=7, which gives most stable ancestry estimates, we observe a partitioning of the KhoeSan ancestry into ‘northern Kalahari’ ancestry shared with Ju|’hoansi and a distinct southern or circum-Kalahari ancestry present in the Nama and the ‡Khomani. On average, in the ‡Khomani San we find 55% northern Kalahari KhoeSan ancestry, 21% southern Kalahari KhoeSan ancestry, 11% European ancestry (common in CEU and French individuals), 12% western African ancestry (common in Yoruba and Bantu-speaking populations), and 2% attributable to other African populations (Tanzanian hunter-gatherers, East African, and North African populations, **Table S2**, Figure 2A and **Figure S2A**). The Nama differ from the ‡Khomani in their proportion of northern versus southern Kalahari ancestry; they have 17% northern Kalahari ancestry, 62% southern Kalahari ancestry, 9% European ancestry, 10% western African ancestry, and 1% attributable to other African populations on average. The western African fraction in the Nama is significantly more variable among individuals (p=1.08e-5), resulting from recent Damara gene flow (Uren et al., 2016). The partition of ancestry components occurs in the same order and is correlated between ADMIXTURE and PCA (Figure 2, Figure S2A,D-F).

**Figure 2.**
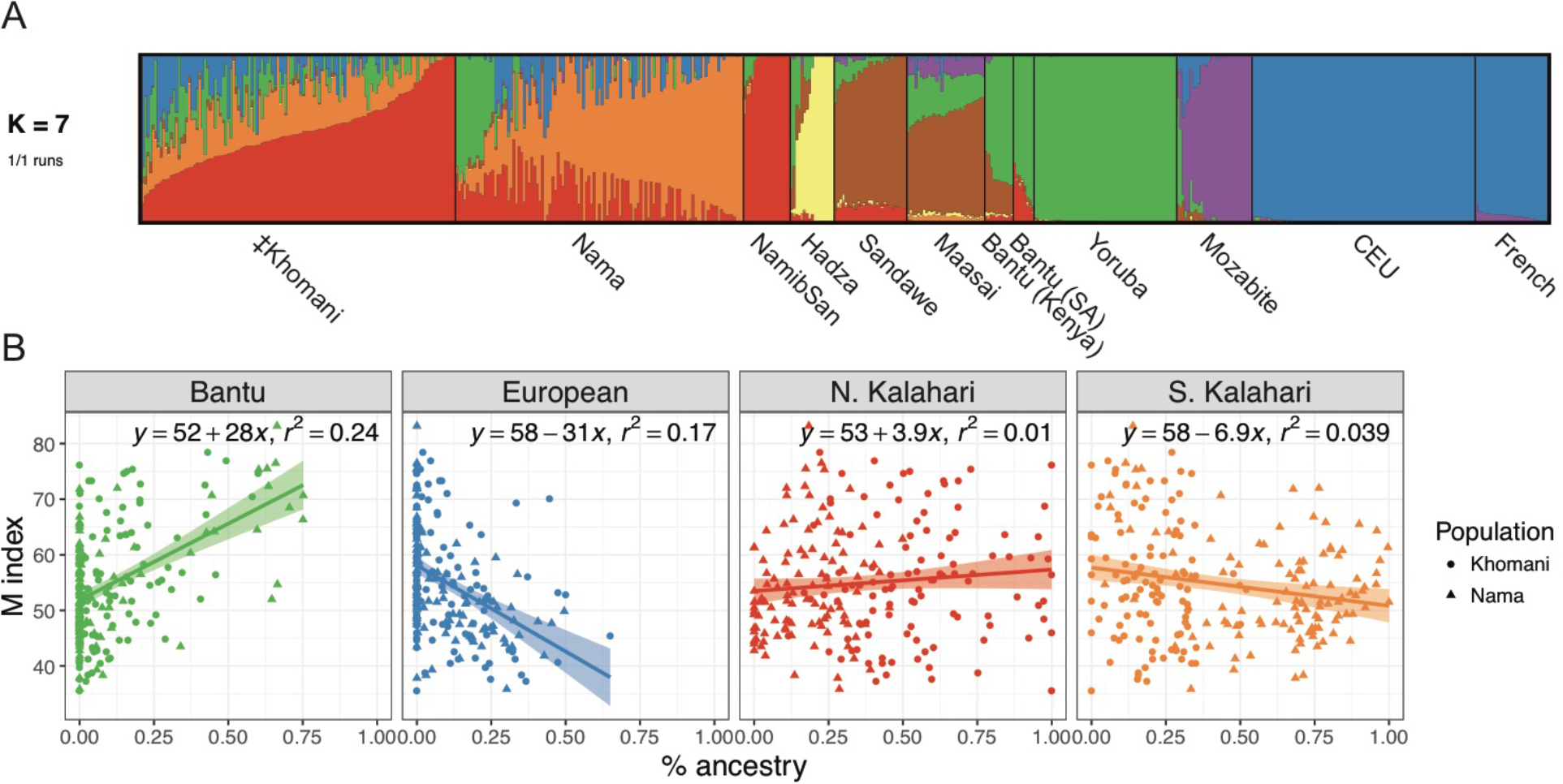
Ancestry components in the KhoeSan and association with pigmentation. **A**) ADMIXTURE proportions at *k*=7 for the ‡Khomani and Nama populations, using Namibian San, Hadza, Sandawe, Maasai, Kenyan Bantu, South African (SA) Bantu, Yoruba, Mozabite, Central Europeans (CEU), and French populations as a reference panel (see also **Figure S2**). **B**) Associations between substantial *k* ancestry clusters and average melanin (M index) baseline pigmentation value in the combined ‡Khomani and Nama populations. The Bantu and European components each constitute ≥ 5% of the total KhoeSan ancestry on average and have significant associations in the best multivariate model(*p* <0.05).

We performed forward stepwise regression to select the best multivariate mixed model of ancestry and pigmentation with a random effect accounting for the genetic relationships among individuals. Sex and age do not significantly correlate with baseline skin pigmentation, suggesting that our quantitative measure of underarm reflectance is not significantly affected by UV exposure. The best model fit, measured via AIC, included Bantu, European, East African, and Hadza ancestries, although the latter two components comprise ≤ 1% of individuals’ total ancestry on average and are likely imprecisely measured. In the multivariate mixed model with European and Bantu admixture components, European ancestry is strongly correlated with lightened skin (β = −18.09, p=2.9e-03), and Bantu ancestry is correlated with darkened skin (β = 25.60, p=1.8e-09). Together, we estimate that fixed admixture effects explained 34% of the variation in skin color (adjusted R^2^); by comparison, 44% of pigmentation variation in Cape Verdeans is explained by admixture effects (Beleza et al., 2013b). Marginal associations are shown in Figure 2B, with pairwise ancestry correlations shown in **Figure S2B**. Southern Kalahari ancestry, frequent in the Nama, is significantly anti-correlated with Bantu ancestry and is marginally predicted to lighten skin, but not when modeled jointly with Bantu ancestry in a multivariate model. Interestingly, the mean pigmentation of Nama and ‡Khomani individuals with <90% KhoeSan ancestry is *not* significantly different from individuals with >90% KhoeSan ancestry (p=0.94), although the variance is significantly greater in more admixed individuals (admixture from either/both European or Bantu ancestries, p=2.2e-3). These results suggest that while admixture increases phenotypic variance, pigmentation alleles on KhoeSan haplotypes contribute more to the overall heterogeneity than those on European or Bantu haplotypes. Consistent with this result, we observe substantial skin pigmentation variation among related individuals, which, coupled with high heritability (see below) suggests a role for large effect sizes of alleles contributing to pigmentation.

### Skin Pigmentation is Highly Heritable

We inferred narrow sense heritability for baseline skin pigmentation and tanning status in the KhoeSan with four methods: family pedigrees 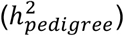, SNP array similarity matrices 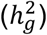, identity-by-descent (IBD) sharing matrices 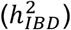, and exome sequence variation (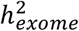, **Table 1**). While pedigree-based heritability estimates are not based on genetic data and therefore not strongly affected by admixture, its careful consideration is necessary for SNP-based estimates, as described previously (Beleza et al., 2013b; Zaitlen et al., 2013; Zaitlen et al., 2014; Thornton et al., 2012) and as we have conducted here. In each of the heritability estimates of baseline skin color, we accounted for admixture proportions with European and Bantu ancestry as covariates, as well as familial relatedness via a kinship covariance matrix. Similarly for tanning status, we accounted for age, sex, and kinship. Previous family-based estimates for skin color heritability in other populations are high, ranging between 55-90% (Byard, 1981; Clark et al., 1981; Frisancho et al., 1981; Harrison and Owen, 1964; Paik et al., 2012). Interestingly, published genetic estimates of skin pigmentation heritability in Europe are low and insignificant, potentially because of reduced genetic diversity at skin pigmentation loci due to positive selection (Zaidi et al., 2017). Our heritability estimates in the KhoeSan are analogous to family-based estimates because of the elevated relatedness in our samples.

**Table 1.** Heritability estimates contrasting baseline skin pigmentation with tanning status. SNP-based heritability estimates were computed with GCTA using genetic relationship matrices (GRMs) calculated from SNP gentoypes, an admixture-corrected GRM computed with REAP, and IBD segments. All models were unconstrained.

| Method | Dataset | SNPs | N | $h^2$ (SE) baseline pigmentation<sup>a</sup> | $h^2$ (SE) tanning status<sup>b</sup> |
| --- | --- | --- | --- | --- | --- |
| GCTA GRM | genotype array | 286,026 | 216 | 0.90 (0.15) | 0.31 (0.19) |
| REAP GRM | genotype array | 286,026 | 216 | 0.97 (0.15) | 0.41 (0.21) |
| K<sub>IBD</sub> | genotype array | NA | 216 | 0.97 (0.16) | 0.45 (0.22) |
| GCTA GRM | exome | 117,132 | 82 | 0.95 (0.26) | 0.37 (0.37) |
| SOLAR | pedigrees | NA | 477 | 0.96 (0.12) | 0.19 (0.11) |

<sup>a</sup> Bantu and European admixture proportions were included as covariates.
<sup>b</sup> Age and sex were included as significant covariates for tanning status (wrist minus baseline underarm pigmentation).

**Table 2.** Replication of previously associated skin pigmentation variants in the joint ‡Khomani and Nama populations. P-value indicates the joint association across all KhoeSan individuals using a linear mixed model accounting for European and Bantu admixture as well as kinship. Beta values reflect the effect size of adding one derived allele, assuming an additive model, to the distribution of M index (see Figure 1).

| Gene | rsID | P-value | Beta | Derived<sup>a</sup><br>frequency | Allele<br>number<sup>b</sup> | San-specific<br>frequency | San 95%<br>CI<sup>c</sup> | W.<br>AFR<sup>d</sup> | N.<br>EUR<sup>e</sup> |
| --- | --- | --- | --- | --- | --- | --- | --- | --- | --- |
| UGT1A | rs6742078 | 0.58 | -0.44 | 0.54 | 460 | 0.60 | [0.54,0.69] | 0.47 | 0.29 |
| SLC45A2 | rs35395 | 0.98 | -0.02 | 0.32 | 882 | 0.21 | [0.18,0.25] | 0.20 | 0.99 |
| SLC45A2 | rs16891982 | 1.2E-03 | -2.84 | 0.14 | 882 | 0.00 | [0.00,0.02] | 0.00 | 0.98 |
| IRF4 | rs12203592 | 0.83 | -0.54 | 0.01 | 882 | 0.00 | [0.00,0.00] | 0.00 | 0.17 |
| IRF4 | rs12202284 | 0.51 | 0.99 | 0.04 | 824 | 0.00 | [0.00,0.01] | 0.15 | 0.21 |
| OPRM1 | rs6917661 | 0.29 | -0.71 | 0.66 | 882 | 0.71 | [0.67,0.79] | 0.61 | 0.76 |
| EGFR | rs12668421 | 0.65 | -0.49 | 0.08 | 882 | 0.02 | [0.01,0.08] | 0.06 | 0.27 |
| TYRP1 | rs13289810 | 0.61 | 0.53 | 0.19 | 882 | 0.18 | [0.11,0.25] | 0.24 | 0.34 |
| BNC2 | rs10756819 | 0.51 | 0.91 | 0.08 | 466 | 0.02 | [0.00,0.05] | 0.07 | 0.65 |
| GATA3 | rs376397 | 0.91 | 0.07 | 0.65 | 872 | 0.79 | [0.75,0.82] | 0.31 | 0.32 |
| GRM5, TYR | rs10831496 | 0.28 | -0.90 | 0.52 | 460 | 0.63 | [0.57,0.70] | 0.12 | 0.69 |
| TYR | rs1042602 | 0.74 | 0.58 | 0.06 | 466 | 0.00 | [0.00,0.02] | 0.00 | 0.38 |
| KITLG | rs12821256 | 0.02 | -5.28 | 0.02 | 882 | 0.00 | [0.00,0.01] | 0.00 | 0.17 |
| OCA2 | rs1800404 | 0.53 | -0.40 | 0.55 | 854 | 0.65 | [0.56,0.74] | 0.11 | 0.81 |
| OCA2 | rs7495174 | 0.92 | -0.07 | 0.71 | 716 | 0.61 | [0.55,0.69] | 0.26 | 0.90 |
| HERC2 | rs12913832 | 0.09 | -1.70 | 0.10 | 882 | 0.00 | [0.00,0.02] | 0.01 | 0.79 |
| APBA2 | rs4424881 | 0.25 | -1.24 | 0.18 | 440 | 0.02 | [0.00,0.06] | 0.07 | 0.86 |
| SLC24A5 | rs1426654 | 9.8E-09 | -3.58 | 0.40 | 882 | 0.24 | [0.17,0.32] | 0.05 | 1.00 |
| MC1R | rs1805007 | 0.80 | -0.64 | 0.01 | 630 | 0.00 | [0.00,0.03] | 0.00 | 0.11 |

<sup>a</sup> Derived alleles were determined previously from great ape genome sequencing (Prado-Martinez et al., 2013).
<sup>b</sup> Allele number indicates the total number of alleles genotyped or sequenced across all KhoeSan samples.
<sup>c</sup> Confidence interval for the San-specific frequencies indicates the allele frequencies specifically on ‡Khomani haplotypes, assessed with local ancestry tracts.
<sup>d</sup> W. AFR (western African) allele frequencies were estimated from 405 ESN, GWD, YRI, and MSL populations in the phase 3 1000 Genomes project
<sup>e</sup> N. EUR (northern Europeans) allele frequencies were estimated from 190 GBR and CEU populations in the phase 3 1000 Genomes project

We first constructed pedigrees from ethnographic interviews for individuals within the ‡Khomani and Nama populations and verified relationships where possible with genetic data. 533 individuals (including parental individuals not sampled) could be assigned to a pedigree, resulting in 354 extended pedigrees and 470 nuclear families. Via traditional pedigree-based estimation of narrow sense heritability using the Sequential Oligogenic Linkage Analysis Routines (SOLAR) software (Almasy and Blangero, 1998), we estimate an 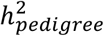 of 0.96 ± 0.12 for baseline skin color. We then asked whether variation present on the ascertained SNP arrays or from exome sequencing could explain a similar fraction of the pigmentation variation. Genetic heritability estimates inferred from recently admixed populations have two potential problems: 1) inferred familial relationships between individuals are less accurate (Thornton et al., 2012), and 2) environmental confounders (e.g. socioeconomic status) could be associated with the variance component attributed to additive genetic effects. In order to address the first issue, we use the proportion of KhoeSan, European and Bantu ancestry per individual to correct the SNP array genetic relatedness matrix (GRM) as described by the REAP approach (Thornton et al., 2012). The REAP matrix is also compared to the IBS matrix inferred using default GCTA parameters that do not account for stratification (*Methods*). We include European and Bantu ancestry as global covariates in the heritability estimation. All further estimation of 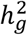 was made using the unconstrained model in GCTA. Furthermore, we contrast baseline pigmentation with tanning status (i.e. sun exposed wrist – underarm melanin pigmentation); if our estimates were inflated by environmental confounders, we would also expect inflated heritability of tanning status.

The array-based heritability point estimates are consistently but not significantly higher when using a kinship matrix from Relatedness Estimation in Admixed Populations (REAP) than GCTA’s identity-by-state (IBS) GRM, both for the joint dataset and each population separately (Table 1 and **Table S3**). We estimate 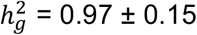 (standard error) in an unconstrained model across both populations using the REAP GRM. We find consistent results from exome sequence data, where we estimate that 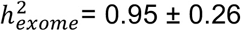 in the ‡Khomani. We then used the familial relationships (**Figure S1**) and population-level endogamy to estimate heritability from IBD sharing among all individuals in the ‡Khomani and Nama; we obtain a similar estimate of 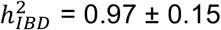(*Methods*, see also (Zaitlen et al., 2013)).

We contrast the high heritability estimates for baseline pigmentation with estimates for tanning status. Tanning status is significantly associated with both sex (male β=6.2 increase in M index, p = 4.2e-4) and age (β=0.18 increase in M index per year, p=1.8e-4), but not with admixture proportions. None of the tanning status h^2^ estimates, including pedigree-, IBD-, exome-, and SNP array-based estimates, are significantly greater than 0 (Table 1), consistent with previous observations that tanning status is largely environmentally determined by UV exposure (Clark et al., 1981; Nan et al., 2009). Previous GWAS of tanning status have also failed to identify and replicate significant SNPs that are not already known to canonically influence baseline pigmentation (Nan et al., 2009). The stark contrast of the baseline pigmentation and tanning status heritability estimates, and the consistency of h^2^ across methods indicates that our high baseline pigmentation heritability estimates do not simply arise from pedigree and population structure and that socioeconomic factors are unlikely to have significant effect on our heritability estimates.

### A Complex Genetic Architecture in the KhoeSan

The genetic architecture of skin pigmentation has been described as simpler than many other phenotypes, for which only a few genes explain ~35% of the total variation in a given population, and average genomic ancestry explains an additional ~44% of the variation, indicating a long tail of smaller effects (Beleza et al., 2013b; Candille et al., 2012). We investigated how much of the heritable variation in KhoeSan populations can be ascribed to previously annotated pigmentation gene sets (Figure 3A). The first gene set (GS1) consists of 14 genes containing or near previously discovered skin pigmentation genetic associations in Europeans, East Asians, Cape Verdeans, and Native Americans (Table 2 and **Table S6**). The larger, second gene set (GS2) contains 50 genes compiled previously (Beleza et al., 2013b) from human pigmentation associations, positive selection scans, and model organism pigmentation loci. The third gene set (GS3) contained 50 loci most significantly associated with pigmentation in the KhoeSan (phase 1, see section titled “*Novel Variants Influence Skin Pigmentation in KhoeSan Populations*”). We partitioned the genome into GS1, GS2, GS3, and the rest of the genome and performed four comparisons, computing the variance explained by: GS1 versus the rest of the genome, GS2 versus the rest of the genome, GS3 versus the rest of the genome, and GS1 versus GS2 versus the rest of the genome. For each comparison, we performed a restricted likelihood ratio test. The GS1 and GS2 gene sets do not explain a significant fraction of the heritability; that is, the heritability estimates overlap with zero. Rather, the remainder of the genome explains the overwhelming majority of the heritability (Figure 3B, σ^2^_GS1_=0.08 vs σ^2^_Genome_=0.82, p_Genome_=2.7e-5; σ^2^_GS2_=0.09 vs σ^2^_Genome_=0.79, p_Genome_=3.3e-4; and σ^2^_GS1_=0.08 vs σ^2^_GS2_=0.09 vs σ^2^_Genome_=0.71, p_Genome_=2.5e-3, respectively). This result contrasts with conclusions from previous studies and indicates that the vast majority of variation in KhoeSan skin pigmentation arises from pigmentation genes yet to be discovered, providing strong evidence for a complex, polygenic architecture. GS3 explains a small but significant fraction of the heritability, as discussed below.

**Figure 3.**
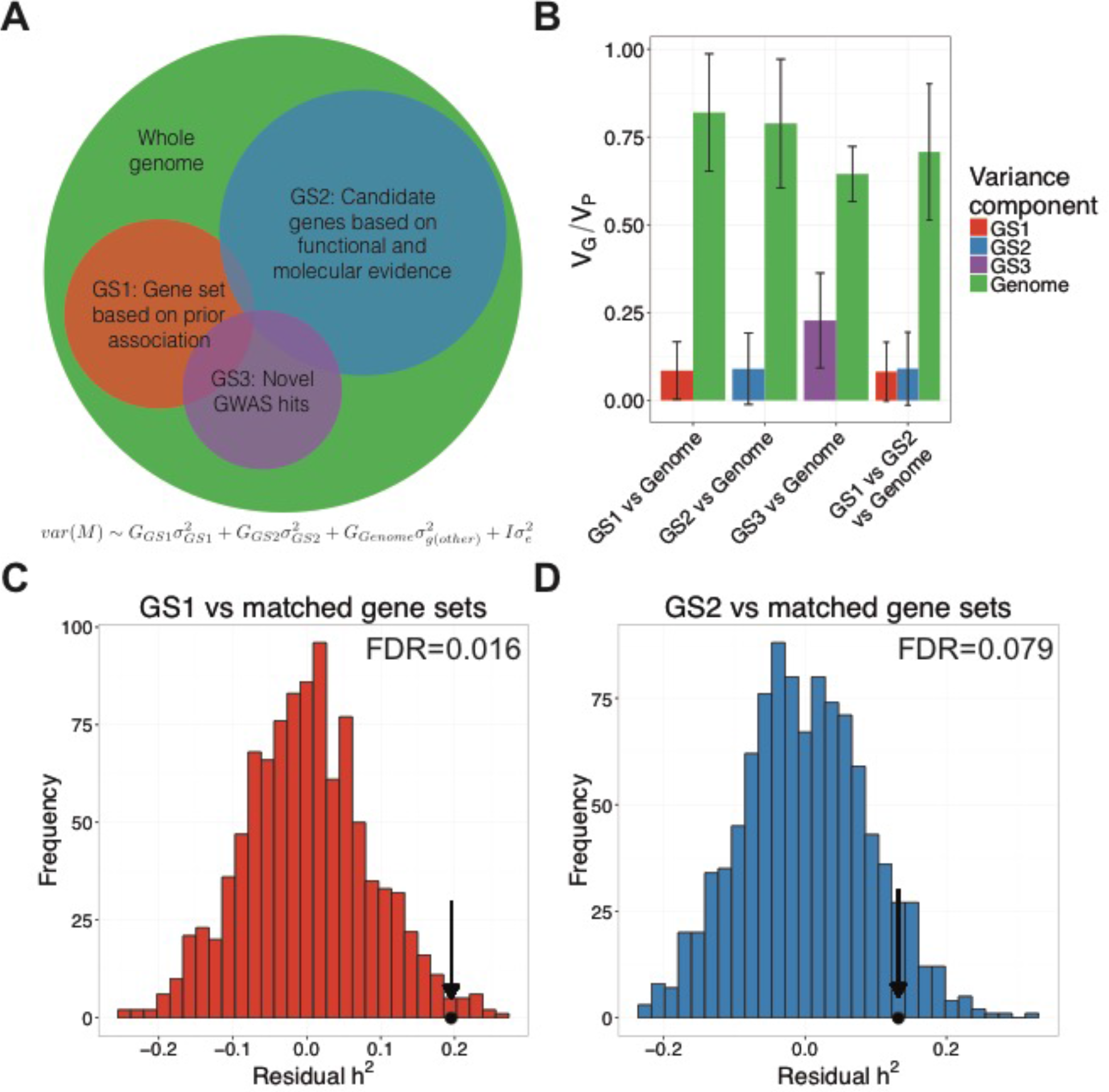
Partitioned heritability across known and novel gene sets. Heritable variation in KhoeSan pigmentation is partially explained by previously associated loci, newly associated loci, and candidate genes discovered in divergence studies of other populations, and in animal models. **A**) Schema illustrating how heritability analyses were used to partition the phenotypic variance explained by candidate gene sets (GS1, GS2) and novel associations (GS3) compared to the rest of the genome. **B**) Variance components analysis in GCTA comparing pigmentation variability explained by GS1, GS2, and the rest of the genome. Error bars span ± 1 standard error. **C**) Heritability explained by estimated value observed in our data (dot and arrow) versus matched null distribution in the ‡Khomani and Nama after accounting for number of SNPs in GS1 gene sets containing 14 genes previously associated with skin pigmentation in other populations. **D**) As in C), where GS2 = gene set from **Table S4** of (Beleza et al., 2013b) compiled based on pigmentation function (see also **Figure S3**).

We further assessed whether GS1 and GS2 explain more of the heritable variation than a random sample of coding regions; genes tend to explain more phenotypic variation than noncoding regions (Gusev et al., 2014). Specifically, we matched both candidate gene sets by number of genes, length, and number of exons and permuted these matched samples 1000 times. After regressing out the effect of variable numbers of SNPs per gene set, we find that both GS1 and GS2 explain more than random genes with a 10% false discovery rate (FDR=0.016 and FDR = 0.079, Figure 3C-D, respectively) across both KhoeSan populations. This is not significant in the Nama alone (**Figure S3**), likely because of ancestry heterogeneity between the two populations.

### Replication of Known Pigmentation Associations in the KhoeSan

Even though previously identified pigmentation loci explain little of the phenotypic variance in our samples, it is possible that these loci simply have small effect sizes in the KhoeSan. We used SNP array and/or resequencing data in a linear mixed model with ancestry covariates (see Methods and “*Novel Variants Influence Skin Pigmentation in KhoeSan Populations*”) to assess both the frequencies and effect sizes of 42 previously identified eye, skin, and hair pigmentation variants, some of which have been experimentally shown to be causal (Table 2 and **Table S6**). To this end, we also deconvolved recent admixture into local ancestry tracts across the genome and estimated the allele frequencies specifically on KhoeSan haplotypes via expectation-maximization (Gravel et al., 2013). Known pigmentation allele frequencies vary considerably between the ‡Khomani San, Europeans, and West Africans (Table 2).

Most previously identified pigmentation associations do not replicate with genome-wide significance or nominally in the ‡Khomani and Nama, with a few exceptions. Four SNPs in the genes *SLC45A2* (rs16891982, p=1.2e-3), *KITLG* (rs12821256, p=0.02), and *SLC24A5* (rs1426654, p=9.8e-9 and rs2470102, p=1.1e-8) marginally replicate in the ‡Khomani + Nama under an additive model. The derived allele frequencies of the associated SNPs in *SLC45A2* and *KITLG* are low in the KhoeSan, consistent with ~10% admixture from recent European gene flow. Interestingly, however, SNPs in *OCA2*, *SLC24A5* and *GRM5/TYR* are at much higher frequencies in both the ‡Khomani and Nama than expected from European admixture alone, as estimated from global ancestry (**Methods**). We do not replicate the vast majority of previously observed skin pigmentation associations in our dataset, potentially due to low frequencies in the KhoeSan, power limitations, differentiated LD structure in which the tag SNPs are non-causal pigmentation alleles, or epistatic effects. It is therefore unsurprising that when we applied forensic models based on only seven SNPs that claim very high prediction accuracy of skin color across populations (>99%) (Spichenok et al., 2011; Hart et al., 2013), we did not find a significant association with quantitatively measured M index (p=0.31, **Figure S5B**).

Because high divergence in a segment of the genome can be a signature of selection (e.g. XP-EHH scans), we assessed genetic divergence between KhoeSan, West African, and European populations at SNPs and in sliding windows across the genome. We find considerable divergence in many canonical pigmentation genes when comparing regions of the genome across populations (Figure 4A-B). We followed up our divergence scan by focusing on two outlier genes that were highly diverged among all three populations: *SLC24A5* and *OCA2* (Figure 4). The divergence in *SLC24A5* is among the highest in the genome, especially between the KhoeSan and European populations (Figure 4D). Interestingly, different regions of *OCA2* exhibit elevated divergence between the KhoeSan and European comparison versus the KhoeSan and West African comparison (Figure 4C). A previous study suggested that the derived, synonymous T allele of rs1800404 in *OCA2* has been positively selected and is a candidate skin pigmentation variant conferring light skin in Europeans and KhoeSan populations based on its global allele frequency distribution (Norton et al., 2007). We confirm its elevated allele frequency on KhoeSan haplotypes (65%), but do not find an association with skin pigmentation (p=0.53). Variants in *OCA2* explain most of the variation in human eye color (Duffy et al., 2007), and rs1800404 was later significantly associated with this phenotype (Eriksson et al., 2010); ‡Khomani and Nama individuals notably have heterogeneous eye color, with a range of brown, hazel, and green eyes. We identified a missense mutation in *OCA2* (rs1800417, ns with skin pigmentation: p=0.87) with a derived allele (G) frequency of 0.32 in the KhoeSan (**Table S6**) that is at low frequency in all other populations surveyed (global allele frequency = 0.016 in Phase 3 1000 Genomes and 0.0058 in the Exome Aggregation Consortium, ExAC).

**Figure 4.**
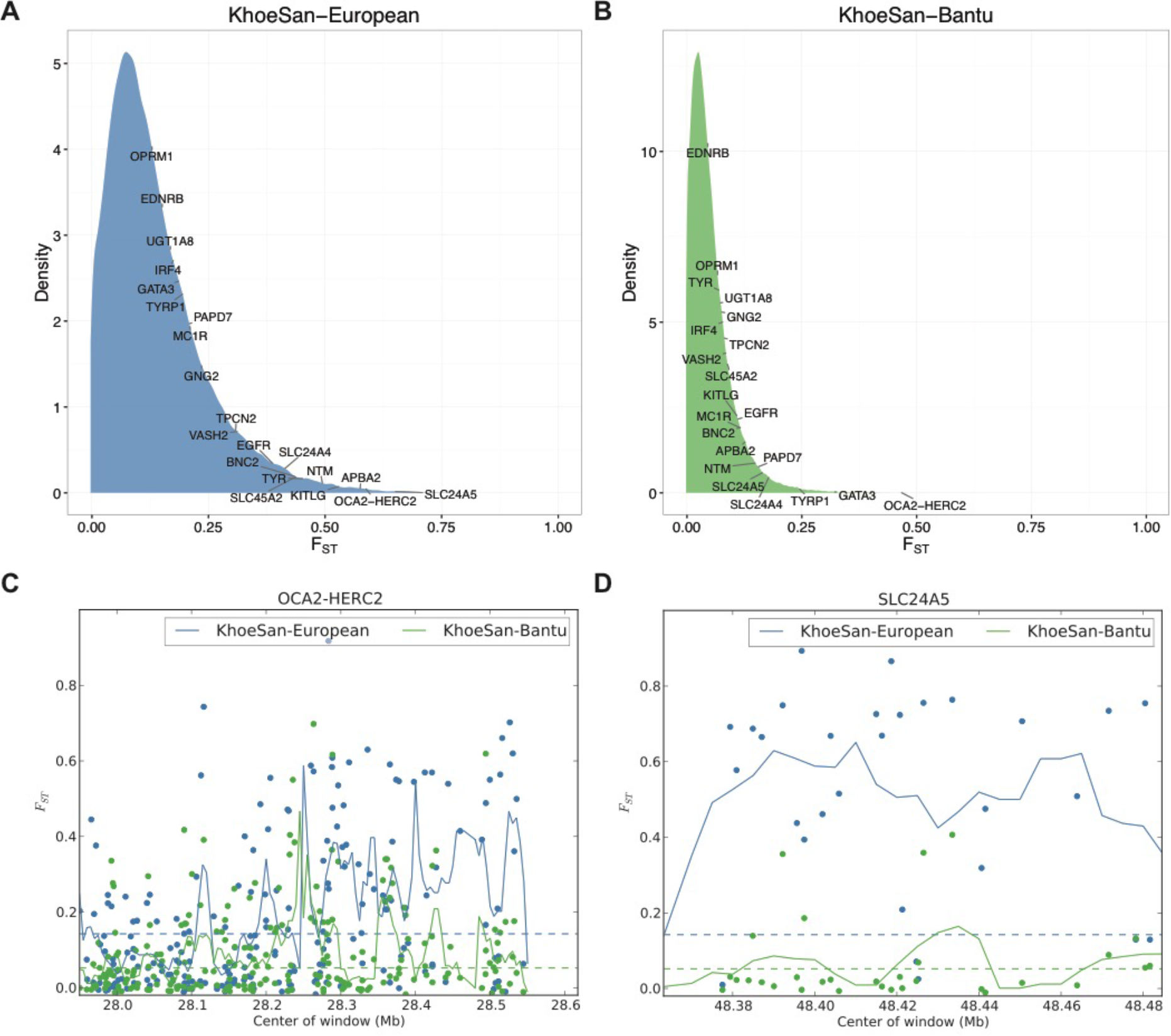
Genetic divergence in genes previously associated with pigmentation. A-B) Distribution of weighted F_ST_ in 20 kb moving windows of SNPs across the genome with a step size of 5 kb. Labels indicate where the maximal F_ST_ window from each canonical pigmentation gene lies in the distribution. Divergence depicted is between A) the KhoeSan and Europeans, and B) the KhoeSan and West African populations. C-D) F_ST_ in canonical pigmentation genes. Dots indicate SNPs, lines indicate moving averages over 20 kb windows with a step size of 5 kb. Canonical pigmentation loci/genes are shown as: C) the *OCA2-HERC2* locus, and D) the *SLC24A5* gene.

### Novel Variants Influence Skin Pigmentation in KhoeSan Populations

To identify novel variants associated with skin pigmentation in the ‡Khomani and Nama, we performed a 2-stage study (**Figure S6A**), employing a linear mixed model approach including recent admixture covariates as fixed effects and covariance matrices adjusted for admixture (akin to a GRM in GCTA) as random effects to identify associations between pigmentation and high quality imputed variants (Alexander et al., 2009). We assessed the quality of the imputation via homozygous reference, heterozygous, and homozygous non-reference concordance with high coverage exome sequencing data (**Figure S4A**). We ran the initial GWAS (i.e. phase 1) with imputed variants from 107 ‡Khomani and 109 Nama individuals (**Table S4**, **Table S5**, **Figure S6A-C**), and the genes closest to the strongest associations (**Table S5**) showed a significant enrichment in multiple mammalian phenotypes related to skin pigmentation (abnormal extracutaneous pigmentation p=2.3e-3, abnormal melanocyte morphology p=5.8e-3, abnormal skin morphology p=3.5e-2). Further, the strongest signals across the genotyped ‡Khomani and Nama cohorts were near canonical pigmentation genes (e.g. *TYRP1* and *SLC24A5*) (Sturm, 2009), genes associated with pigmentation-related disorders (e.g. *TYRP1*) (Jin et al., 2010), or genes implicated in pigmentation in model organisms and *in vitro* studies (e.g. *VLDLR*, *SMARCA2*, and others) (Demir et al., 2013; Keenen et al., 2010; de la Serna et al., 2006; Liu-Smith et al., 2014). To assess the variation explained by the most significantly associated loci, we generated an additional gene set, referred to as “GS3”, using the 50 most significantly associated loci ± 10 kb. We find that the GS3 loci explain significantly more of the heritable variation in skin pigmentation than previously identified pigmentation candidate genes in the KhoeSan, but that the majority of heritable variation remains to be explained (Figure 3B, σ^2^_GS3_=0.23±0.13, p_GS3_=0.027 vs σ^2^_Genome_=0.64±0.08, p_Genome_<1e-5).

Based on initial evidence from the imputed ‡Khomani pigmentation GWAS, we designed a targeted NGS capture and successfully resequenced 36 candidate pigmentation regions (**Table S7**, **Figure S6**) across a larger set of 451 KhoeSan samples in order to improve power to detect associated loci (**Table S8**, Supplementary Materials), including 269 Khomani, and 182 Nama individuals. In this larger sample, we observe more variants significantly associated with pigmentation than expected by chance in the resequencing regions (Figure 5A). The strongest signal comes from SNPs in *SLC24A5*, 8 of which are all in high pairwise LD (R^2^ > 0.6) on a high frequency haplotype (Figure 5B). We identify significant associations between lighter skin and derived *SLC24A5* SNPs, including the putatively causal p.Thr111Ala rs1426654 allele (β=−3.58 on M index scale, p=9.8e-9), which has previously been associated with skin pigmentation in Eurasians. The most strongly associated SNP (rs2555364, β=−3.58 on M index scale, p=6.7e-9) is tightly linked with rs1426654 (LD R^2^=0.81). These variants are strongly differentiated between Europeans and Africans, with rs1426654 having derived allele frequencies of 99.7% vs 5.5% in 1000 Genomes (excluding ASW and ACB populations with recent European admixture), respectively. The derived allele of rs1426654 has previously been observed in HGDP Ju|’hoansi San samples at 7% frequency, which have no detectable recent European admixture (Norton et al., 2007). The frequency of the derived rs1426654 allele is 40% in the combined Nama + ‡Khomani dataset, which is significantly greater than expected from ~11% European admixture alone (binomial test p=7.8e-52, **Table S6, STAR Methods**).

**Figure 5.**
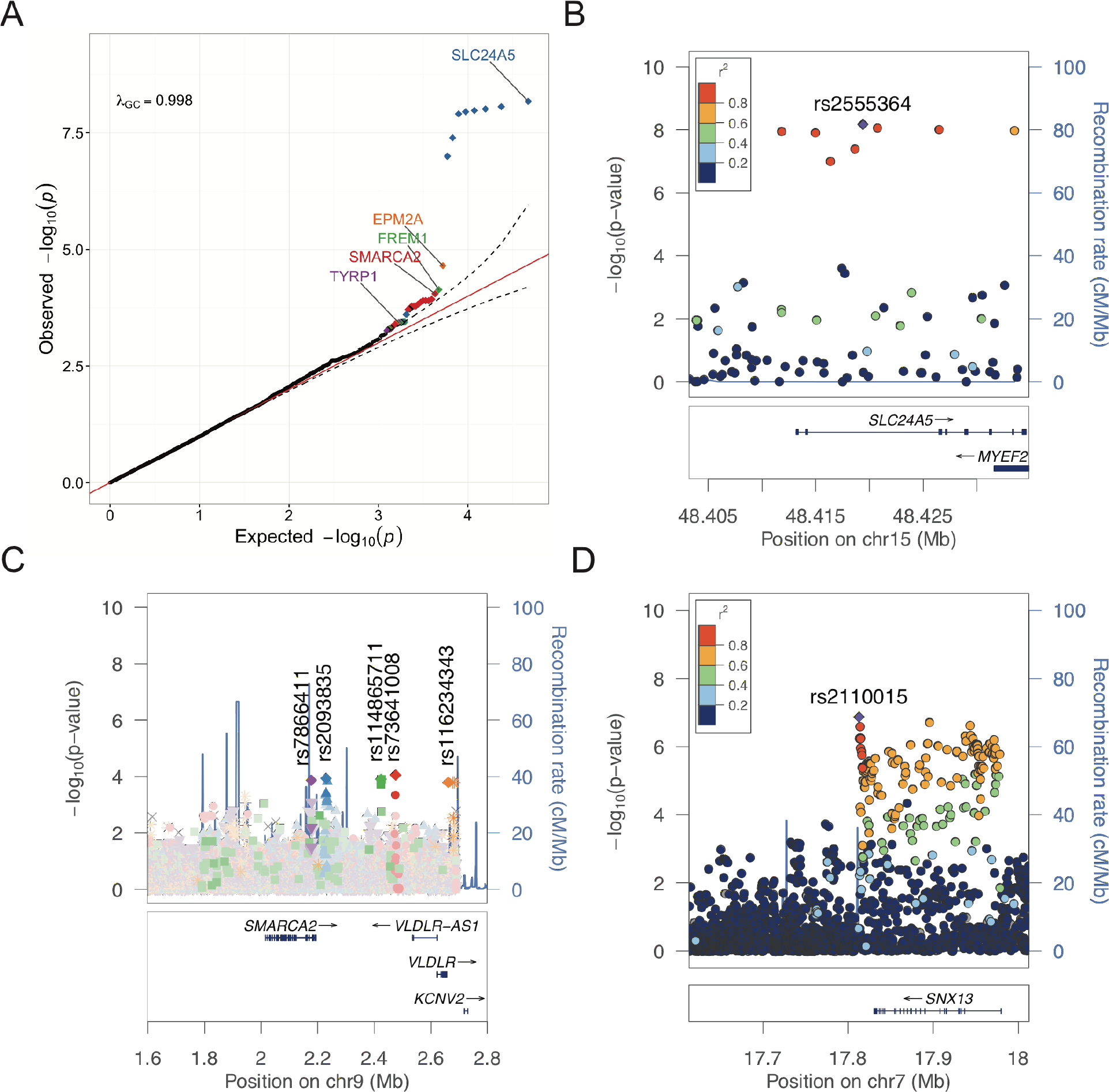
Associations between genetic data and baseline pigmentation. Information regarding targeted resequencing regions is shown in **Table S7**. A) Targeted resequencing QQ plot. 95% confidence interval on the QQ plot is drawn assuming the *j*^th^ order statistic from a uniform sample follows a *Beta*(*j, n – j* + 1) distribution. Colors differentiate loci containing more than one variant associated more significantly than the 95% confidence interval in a region. B-C) LocusZoom plots of targeted resequencing genetic associations incorporating KhoeSan-specific LD. Recombination rates are from HapMap b37. Regions include: B) SLC24A5, and C) 5 independent signals associated with p < 1e-3 in/near SMARCA2 and VLDLR. E) LocusZoom plot of suggestive association in/near SNX13 from meta-analysis of phase 1 and phase2 (**Figure S7A**) imputed associations with KhoeSan-specific LD.

Multiple low frequency (<5%) SNPs near several additional genes, including *EPM2A*, *FREM1*, *SMARCA2/VLDLR*, and *TYRP1*, are above the 95% confidence interval of expected versus observed significance (Figure 5). Two of these regions are near *EPM2A* and *FREM1*, which are known to play roles in myoclonic epilepsy and the development of multiple organ systems, respectively; however, neither of these genes play any known role in skin pigmentation either in humans or model organisms. In contrast, there are >5 independent low frequency signals upstream, downstream, and in introns of *SMARCA2* and near *VLDLR* with p<1e-3, with rs7866411 (p=8.91e-5) and rs2093835 (p=1.17e-4) being the SNPS most significantly associated with skin pigmentation. We used HaploReg to infer regulatory activity in/near these peaks and identify multiple enhancer and DNAse peaks identified in skin, including melanocytes and/or keratinocytes, overlapping top tag and/or perfectly linked SNPs (**Table S9**). We also identify a low frequency association (rs34803545, p=3.7e-4) ~600 kb upstream of *TYRP1* in a gene desert. This variant is perfectly linked with multiple conserved variants, one of which exhibits enhancer activity and DNAse hypersensitivity specifically in skin (**Table S9**).

We followed our phase 1 GWAS analysis with a 2^nd^ phase, in which an additional 240 unique individuals were genotyped (**Table S4, Figure S6A**) and meta-analyzed with phase 1 summary statistics. While two tanning status associations met genome-wide significance, none of the loci contained linkage peaks, suggesting that they are most likely spurious. The tanning status GWAS results are expected from a phenotype with low heritability. As expected from the resequencing study, we identified a genome-wide significant association in *SLC24A5* (rs2470102 derived allele β =-3.4, p=3.6e-12) and a suggestive association upstream of *TYRP1* (chr9:12088112, frequency=0.014, β = −13.6, p=1.1e-07 **Figure S6B-C**, **Figure S6F-G**). We identified an additional suggestive novel association in and near *SNX13*, with common derived T alleles of rs2110015 associated with light skin (β = −3.1, p=1.3e-07, **Figure S6H**); *SNX13* regulates lysosomal degradation and G-protein signaling, but has not previously been associated with skin pigmentation.

## Discussion

Pigmentation has been described previously as a relatively simple trait with few loci of large effect contributing to the phenotype (Hart et al., 2013; Spichenok et al., 2011; Sulem et al., 2007). However, populations living in continental Africa, where humans have the greatest genetic diversity and variation in pigmentation (as demonstrated here), have been largely ignored in genetic studies of quantitatively phenotyped pigmentation. We investigated the genetic architecture of pigmentation in two KhoeSan populations: the Khomani San and Nama, where baseline melanin variation is substantial. The southern African KhoeSan populations are the most polymorphic modern human populations yet studied (Henn et al., 2011), and provide a unique glimpse into the evolution of pigmentation.

### Novel Genetic Associations with Pigmentation

We have performed the first genetic discovery effort for pigmentation loci in the Nama and ‡Khomani San populations. The strongest allelic associations include previously associated variants, noncoding regions near canonical pigmentation genes, and novel genes shown in model organisms to have a role in pigmentation. The strongest association is in *SLC24A5*, which is a well-known pigmentation gene (Lamason et al., 2005) and is among the most differentiated regions of the genome between European and African populations – indicative of strong positive selection in northern Europeans (Sturm and Duffy, 2012). We find that derived variants in *SLC24A5* are at high frequency in the KhoeSan, including missense mutations that influence skin and eye pigmentation (Table 2). Notably, these variants are segregating at higher frequency than expected by recent European admixture alone. Three possible evolutionary scenarios that may explain these elevated frequencies are: 1) these variants arose in southern Africa more than 100,000 years ago and were later selected for in Europeans after the out-of-Africa migration in response to northern UVR environments. Alternatively, 2) these variants arose in Europe/Near East, were introduced into KhoeSan populations via “back to Africa” migration into southern Africa predating 17^th^ century European colonialism (Tishkoff et al., 2007; Pickrell et al., 2012; Pickrell et al., 2014; Uren et al., 2016), and have since been positively selected in the KhoeSan. Lastly, 3) a recurrent mutation (G to A transition at the CpG ancestral dinucleotide, a class of mutations shown to have elevated mutation rates) occurred. Considerable future work is needed to definitively disentangle these scenarios.

We find a significant enrichment of genes related to melanogenesis in our GWAS. Specifically, we find several independent associations near *SMARCA2* and *VLDLR. SMARCA2* has a known role in folate biosynthesis, in vitamin D-coupled transcription regulation, and is differentially expressed across CEU and YRI populations in lymphoblastoid cell lines (Duan et al., 2009).

Additionally, previous functional studies have shown that *MITF*, the transcription factor known as the “master regulator of melanogenesis” due to its ability to activate many melanocyte-specific genes (Praetorius et al., 2013), recruits critical components of the SWI/SNF chromatin remodeling complex, including *SMARCA2*, to the promoter region of its targets (Vachtenheim et al., 2010). This recruitment is required for normal expression of many *MITF* target genes, including *TYR, TYRP1, DCT, RAB27A, BCL2*, among others (Keenen et al., 2010; de la Serna et al., 2006). Additionally, *VLDLR* knockout mice exhibit hypopigmented retinas (Xia et al., 2013). We also find a suggestive association upstream of *TYRP1* (Figure 5A). *TYRP1* mutations in humans have been associated with oculocutaneous albinism and shown to cause nearly Mendelian inheritance of blond hair in Solomon Islanders (Kenny et al., 2012; Sarangarajan and Boissy, 2001). Thus, we observe enrichments of molecular pathways involved in pigmentation beyond those previously identified as associated with the phenotype in non-African populations.

### The Polygenic Architecture of Pigmentation in Africa

We assessed the heritability of baseline skin pigmentation, and find that it is virtually completely heritable in our KhoeSan sample. In contrast, tanning status is primarily environmental, with heritability estimates which are not significantly different from zero. In European populations, predictive models based on only 9 SNPs capture up to 16% of the variance in skin pigmentation (Liu et al., 2015), highlighting its relative simplicity. We applied a predictive model (Spichenok et al., 2011; Hart et al., 2013) based on these SNPs to the Nama and ‡Khomani San populations, and find no significant association between predicted skin color and spectrophotometrically measured skin M index, showing that this estimation fails to capture the genetic variation driving the phenotype in the KhoeSan. Given the large effect sizes and high fraction of variation explained in Eurasian populations, we asked whether and how much of the phenotypic variation can be explained by previously identified genes. All gene sets, including previously associated loci, canonical pigmentation genes, and the most significantly associated variants in this study, explained a small fraction of the phenotypic variance (σ^2^_GS1_=0.08, σ^2^_GS2_=0.09, σ^2^_GS3_=0.23, respectively). As expected from previous work (Martin et al., 2017), our results indicate that genetic risk prediction is strongly affected by population structure. Most of the pigmentation variability in KhoeSan populations is not explained by previously identified loci, suggesting that more than 50 loci (and indeed, likely far more, given our genomic heritability estimates) with a distribution of mostly small effects contribute to variation in pigmentation in the KhoeSan. This suggests that skin pigmentation is a far more complex trait than previously discussed, analogous to numerous other complex traits discussed in biomedical literature.

### The Evolution of Skin Pigmentation: Selection and Constraint

By aggregating a large set of quantitative skin pigmentation phenotypes (N=4,712) from globally diverse populations, we have demonstrated heteroskedasticity as a function of latitude. As observed previously, we find a strong correlation between absolute latitude and average skin pigmentation reflectance caused by melanin content. We also observe that populations with lighter skin have reduced variation within any given study: populations furthest from the equator have narrower distributions, while populations closest to the equator have wider distributions. These patterns suggest that selection is acting differently at different latitudes. In equatorial regions, strong directional selection for darker pigmentation has shifted the distribution means in some populations to M indices greater than 90, but with wide variances. This is consistent with a ‘threshold’ model (Chaplin, 2004) in which the protective benefit of melanin needs to meet some minimum threshold but with no penalty to darker pigmentation; alternatively, diversifying selection could maintain the wide variance.

In stark contrast, pigmentation in far northern European and Asian populations has been under directional selection for decreased melanin production, reflected by very narrow distributions. There may be biological constraints on the lower boundary of skin pigmentation, and/or due to the strong positive selection acting on a few large-effect alleles, there is little genetic variability left at these pigmentation loci. This would simplify the genomic architecture, with relatively few alleles of large effect driving the phenotype, particularly alleles that lighten skin at extreme northern latitudes, and could explain why prior investigations observed an almost Mendelian inheritance of large effect light pigmentation alleles.

Finally, populations at intermediate latitudes have increased variance and higher means than populations in northern Eurasia, but less than equatorial populations. The most parsimonious explanation for this pattern is that stabilizing selection affects the light and dark tails of the pigmentation distribution (Barton, 1999). The Nama and ‡Khomani San appear to be two such instances of this intermediate variation within Africa, likely attributable to their geographic distance from the equator in far southern Africa (~24-29 degrees South). The observed mean and variance differences across the full spectrum of skin pigmentation by latitude may be driven by imbalanced opposing adaptive pressures, where selective forces to produce vitamin D and protect folate from photolysis are unequal and change in response to UV radiation exposure. Given our heritability results and the observed variability in baseline pigmentation; light skin pigmentation in the KhoeSan appears to be due to a combination of many small-effect mutations as well as some large-effect variants. The evolution of the pigmentation phenotype in these populations cannot be explained in terms of only a few variants segregating in Eurasians. A fuller characterization of the genes underlying the architecture in Africans is needed before we can distinguish between the hypothesis of directional versus stabilizing selection across different latitudes (Berg and Coop, 2014).

### Conclusion

Because African populations often carry the ancestral (i.e. dark) allele for skin pigmentation genes identified in Eurasians, allusions to African skin pigmentation have ignored the great variability in this phenotype across Africa. Here, we reiterate that skin pigmentation varies more in Africa than any other continent, and we show that pigmentation in African populations cannot simply be explained by the small number of large effect alleles discovered in Eurasians. Even in light to moderately pigmented KhoeSan populations, the polygenicity of skin pigmentation is much greater than Eurasians, encompassing both known pigmentation genes as well as novel loci. We argue that the distributions of skin pigmentation globally suggest different forces of selection operating at various latitudes. To better understand baseline pigmentation, one of the most rapidly-evolving traits and strongest cases for positive selection in humans, it is essential to quantitatively measure and study pigmentation in a large set of genetically diverged populations that have historically been exposed to different levels of UV radiation. As human genetics moves to ever larger studies of complex traits (Wood et al., 2014), the full picture of genetic architecture will remain incomplete without representation from diverse worldwide populations.

## Author Contributions

BMH, CRG, MWF, and CDB conceived of and designed the study. ARM and AS designed experiments. ARM, MS and AS performed experiments. ARM, JMG, ML, EGA and XL analyzed data. BMH, CRG, CJW, JWM, ARM, JMG, and MM collected samples and measured skin photometrics. EGH, CRG, DMK, MWF, CDB, and BMH supervised the study. ARM and BMH wrote the manuscript with input from ML, JMG, MM, DMK, EGH, MWF, and CRG. All authors read and approved of the final version of the manuscript.

## Acknowledgments

We thank the Beijing Genome Institute and Agilent Technologies for sequencing support. We thank George Chaplin and Nina Jablonski for providing summary statistics for baseline pigmentation in the Xhosa and Cape Coloured populations. We thank Sandra Beleza, Heather Norton, Esteban Parra, Chandana Basu Mallick, Andres Moreno-Estrada and Karla Sandoval, Richard Cooper, Sophie Candille, and their colleagues for providing quantitative phenotypes from their pigmentation work. We thank Catherine Guenther for discussing molecular strategies for functional followup. We would also like to thank the Working Group of Indigenous Minorities in Southern Africa (WIMSA), community leaders and the South African San Institute (SASI) for their advice and oversight. Finally, we thank Richard Jacobs, Wilhelmina Mondzinger, Hans Padmaker, Willem de Klerk, Hendrik Kaiman, and the communities in which we have sampled; without their support, this study would not have been possible. Funding for this project was provided by a Stanford University CDEHA seed grant (NIH, NIA P30 AG017253-12) to BMH and by a trainee research grant to ARM by the Stanford Center for Computational, Evolutionary, and Human Genomics. Funding was also provided by the Morrison Institute for Population Studies, Stanford University. Funding for genotyping additional KhoeSan individuals was provided by the Stanford Center for Computational, Evolutionary, and Human Genomics. ARM was funded by NIH Genetics and Developmental Biology Training Program T32 GM07790. EA was supported by an NIH IRACDA fellowship to Stony Brook University. CRG was supported by the Stanford Genome Training Program 5T32HG000044. CJW, MM and EGH are partly supported by the South African Medical Research Council. The content is solely the responsibility of the authors and does not necessarily represent the official views of the South African Medical Research Council.

## Conflicts of Interest

CRG and BMH own stock in 23andMe, Inc. CRG is a member of the scientific advisory board and academic founder for Encompass Bioscience, Inc. JAG is an employee of AncestryDNA. CDB is a member of the scientific advisory board for Liberty Biosecurity, Personalis, Inc., 23andMe Roots into the Future, Ancestry.com, IdentifyGenomics, LLC, Etalon, Inc., and is a founder of CDB Consulting, LTD. All other authors declare that they have no competing interests.

§ We use the term “KhoeSan” to refer to a diverse array of indigenous populations in southern Africa that carry KhoeSan ancestry and speak Khoe, !Ui-Tuu or Kx’a languages. “KhoeSan” is not accepted by all such communities; where possible we refer to populations by their specific ethnic name. This grouping lumps together populations of different languages, cultures and variable genetic diversity.

